# Unravelling the Core and Accessory Genome Diversity of *Enterobacteriaceae* in Carbon Metabolism

**DOI:** 10.1101/2025.02.12.637621

**Authors:** Nicolas Näpflin, Christopher Schubert, Lukas Malfertheiner, Wolf-Dietrich Hardt, Christian von Mering

## Abstract

*Enterobacteriaceae* are commonly colonizing animal guts and impact health. While strain-specific metabolic features can promote gut colonization, we lack systematic knowledge regarding metabolic diversity and the core metabolism shared among *Enterobacteriaceae*. To address this gap, we have analyzed the pan-genome of nearly 20,000 genomes. We found that genes necessary for monosaccharide-fuelled mixed acid fermentation are part of the *Enterobacteriaceae* core genome, while most genes for anaerobic respiration and most carbohydrate utilization genes belong to the accessory genome. Understanding *Enterobacteriaceae’s* metabolic capacity helps clarify the distinction of nutrients consumed by all *Enterobacteriaceae*, and niche-defining nutrient sources, which are genus-, species- or strain-specific. This knowledge sheds light on bacterial nutrient exploitation during gut colonization in health and disease, aiding in the development of targeted interventions for microbiome research and infectious disease control. The theoretical framework described here can also be adapted to analyze core physiological characteristics of other microbiota taxa.

## Introduction

The *Enterobacteriaceae* family consists of diverse Gram-negative bacteria, including both harmful pathogens and commensal members. This family plays a critical role in public health and veterinary medicine, as it is responsible for a large number of foodborne infections worldwide ^1–4^. A defining trait of *Enterobacteriaceae* is their facultative anaerobic lifestyle, allowing them to thrive with or without oxygen. They play an important role in early neonatal life by consuming oxygen in the gut, thereby creating beneficial conditions for obligate anaerobe colonization ^5, 6^. On the other hand, *Enterobacteriaceae* can also play a prominent role in disease, e.g., in inflammatory bowel disease ^7, 8^ or in diarrheal infections ^4^. Extensive research has focused on the virulence factors of pathogenic *Enterobacteriaceae* within genera such as *Escherichia*, *Salmonella*, *Citrobacter*, and *Shigella*, which can cause disease in a wide range of mammals ^9–12^.

Growth in the animal gut lumen is important for commensal and pathogenic *Enterobacteriaceae*, alike. However, it is still not fully understood how *Enterobacteriaceae* strains can establish themselves in a gut which has already been colonized by another related strain. Rolf Freter’s concept of nutrient niches posits that two related bacterial strains can only co-exist if each occupies a preferred niche based on its superior use of a specific nutrient in a well-mixed environment ^13, 14^. This theory extends to spatial structuring within the gut, known as the Restaurant Hypothesis ^15–18^. Nutrient competition among microbiota is a key factor in colonization resistance, where the gut microbiota helps protect against invading bacteria. The central principle is metabolic resource overlap, where microbes with overlapping metabolic needs compete for the same nutrients, effectively blocking access for others. This nutrient competition, which is shaped by the diversity of the microbial community, is particularly intense when the microbiota comprises members closely related to the invading pathogen ^19, 20^. However, our understanding of how widely metabolic characteristics are shared among the *Enterobacteriaceae* family remains limited.

Pan-genome analysis assesses the entire genomic repertoire of a bacterial clade. It encompasses the core genome, which consists of genes present in most genomes, and the accessory genome, which includes lineage-specific genes and unique genes (singletons), which may often be acquired through horizontal gene transfer. The accessory genome is a key source of functional and genetic diversity ^21^. Many bacterial species exhibit an open pan-genome, which tends to expand as more genomes are analyzed ^22, 23^. This underscores the need for comprehensive datasets to capture the full extent of genomic diversity. Advancements in high-throughput sequencing have led to an increase in available genomes ^24, 25^. Tools for pan-genome analysis vary in their approaches. For closely related species, whole-genome alignment strategies can be used ^26^, while larger and more diverse pan-genome analyses rely on protein clustering and e-value cut-offs, bit score ratios, or sequence identity thresholds to define gene families ^27–32^. Challenges such as contamination, fragmented assemblies, and especially computational complexity due to all-against-all comparisons can hinder the analysis of large datasets ^33, 34^. Tools like Roary and PIRATE address these issues by first clustering highly similar sequences to facilitate the analysis of thousands of genomes. Despite these advancements, a comprehensive analysis of the *Enterobacteriaceae* pan-genome is still lacking.

Previous studies on bacterial pan-genomes have predominantly highlighted differences rather than commonalities, often concentrating on species-level distinctions. We utilized an innovative bioinformatics approach to categorize core and accessory genes across an extensive dataset of *Enterobacteriaceae* genomes. Our analysis covered sixteen *Enterobacteriaceae* genera, such as *Salmonella*, *Escherichia*, *Klebsiella*, *Yersinia*, *Enterobacter*, and *Shigella*. Furthermore, our approach aims to uncover similarities across *Enterobacteriaceae*, particularly in energy metabolism, while the dataset provided here can also be used to study other aspects of *Enterobacteriaceae* physiology. Additionally, future studies can build on this workflow to assess conserved features of the metabolism of other bacterial taxa, including pathogens and key microbiota members like *Bacteroidota* and *Firmicutes*.

## Results

### Bioinformatics workflow for analysing the pan-genome of *Enterobacteriaceae*

We analyzed the *Enterobacteriaceae* pan-genome across 16 genera and 80 species, encompassing nearly 20,000 genomes (**Supplementary Table S1**). A brief overview of the bioinformatic workflow is shown in **Fig. 1a**. High-quality genomes (completeness ≥ 95%, contamination ≤ 5%) were selected from progenomes3, and species represented by fewer than 10 genomes were excluded. Overrepresented species (mainly *Escherichia coli*, *Klebsiella pneumoniae* and *Salmonella enterica*) were subsampled randomly (**Fig. 1b**). Orthologous gene families were identified over a range of amino acid identity thresholds from 50% to 95% (in increments of 5%, see Methods). The pan-genome includes nearly 95 million gene sequences clustered into 141,675 gene families. Gene ubiquity follows a universal U-shape curve ^35^, with 1.2% (1,801) of gene families being part of the core genome (present in at least 95% of genomes) (**Fig. 1c**). We found that almost 60% of gene families are unique to specific genera, from which 48% are unique to a single genome (singletons), highlighting high intra-family genetic diversity (**Fig. 1d**). Gene families were annotated against the UniprotKB/Swiss-Prot database ^36^, and categorized into three types: Resolved (i.e., 90% or more sequences with the same annotation), Ambiguous (mixed annotations), and Non-annotated (**Fig. 1a**). The more ubiquitous gene families tend to be well-annotated, while rare gene families (present in less than 5% of genomes) make up over 90% of the pan-genome and remain largely unannotated. Finally, we annotated all genes with Clusters of Orthologous Groups (COG) functional categories using eggnog-mapper (**Fig. 1e**)^37^. Translation, amino acid metabolism and cell wall/ membrane biogenesis represent the largest COG categories within the core genome. On the contrary, the “*function unknown*” category is by far the largest COG category within the accessory genome.

**Figure 1.**
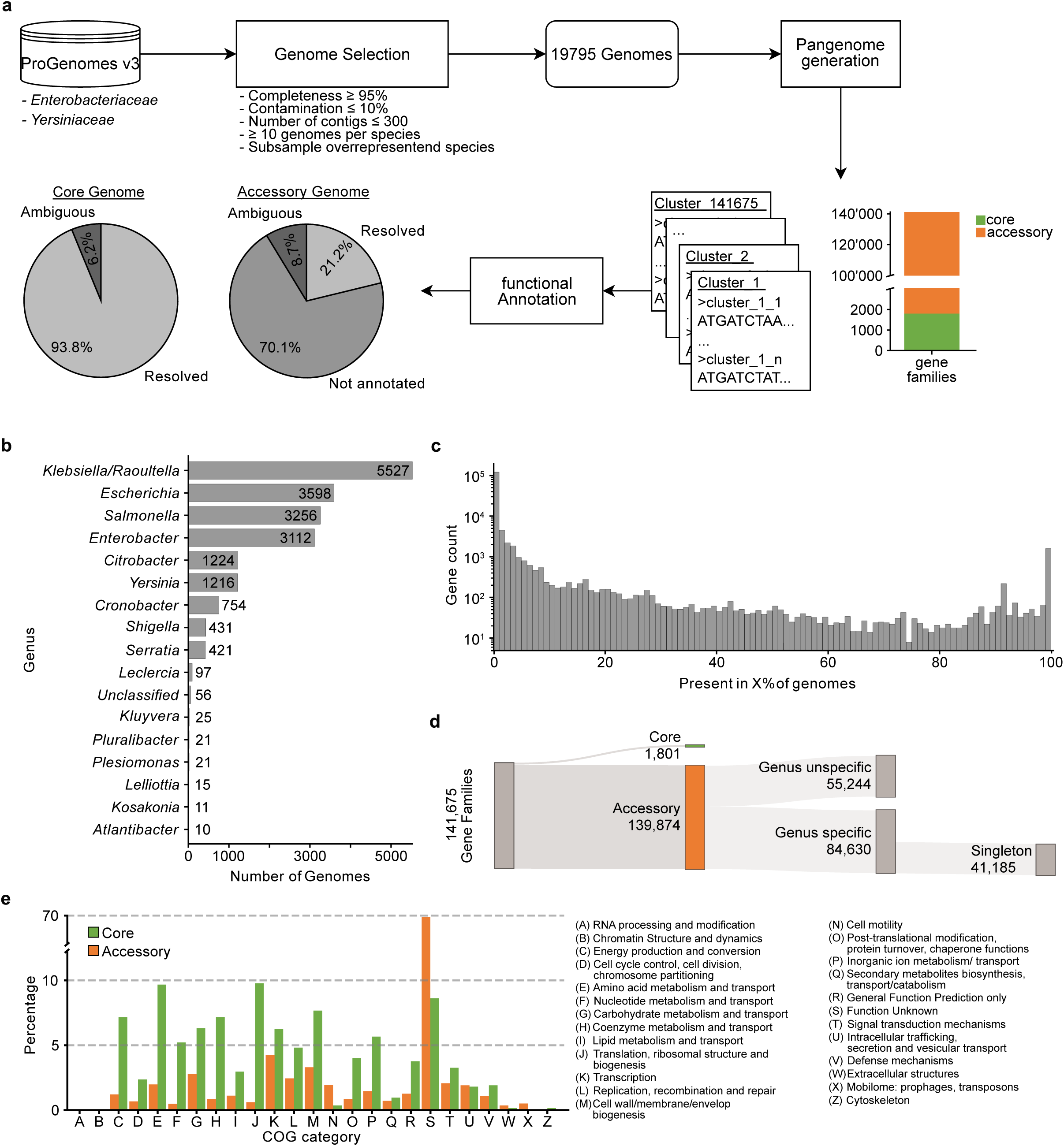
Overview of the bioinformatics workflow. **a** The *Enterobacteriaceae* pan-genome currently includes 141’675 gene families. Gene families are grouped based on their annotation status: Resolved (>= 90% of the sequences contain the same annotation), ambiguous (< 90% of the sequences contain the same annotation and < 10% unknown), and not annotated (>= 10% of the sequences are not annotated). **b** A total of 19’795 *Enterobacteriaceae* genomes across 16 genera were analyzed. **c** Gene ubiquity distribution (expressed as fraction of genomes containing a gene) following a U-shaped curve. **d** Distribution of gene families into core and accessory genome. The accessory genome is further split into genus specific and unspecific families. Almost 50% of genus specific gene families are unique to a single genome (singleton). **e** The functional categories of the accessory genome are dominated by gene families with unknown functions (68.2%). Most frequent function in the core gene families are Translation (9.8%), Amino acid metabolism and transport (9.7%) and cell wall/membrane/envelope biogenesis (7.7%).

### Phylogenetic analysis of the *Enterobacteriaceae* genomes

The *Enterobacteriaceae* family, the largest within the *Enterobacterales* order, is closely related to *Vibrionaceae* and *Pasteurellaceae*. We performed phylogenetic analysis using 103 concatenated core proteins (**Fig. 2, Supplementary Table S2**). It supports the monophyly of many genera within *Enterobacteriaceae* ^38^ and aligns with the 16S rRNA-based phylogenetic tree from the All Species Living Tree project ^39^. Historical classification grouped *Enterobacteriaceae* into core and peripheral members based on DNA-DNA hybridization in relation to *Escherichia coli*. Core members include *Enterobacter*, *Klebsiella*, *Citrobacter*, and *Salmonella*, which are more closely related to *Escherichia coli* in the core genome tree ^40^. The close relationship among *Escherichia*, *Citrobacter*, and *Salmonella* has been confirmed by various single-gene and 16S rRNA-based studies ^38, 41–43^. In contrast, genera such as *Plesiomonas*, *Yersinia*, and *Serratia* appear more peripheral. *Plesiomonas*, initially thought to belong to *Vibrionaceae* based on 5S rRNA, is now placed within *Enterobacteriaceae* due to 16S rRNA and multilocus sequence typing, though its position remains debated ^44^. The placement of *Yersinia* and *Serratia* within *Enterobacteriaceae* is less debatable and consistent with previous studies ^40, 45^. Notably, *Shigella* genomes are mixed within the *Escherichia* clade and do not form a distinct monophyletic group, supporting the view that *Escherichia* and *Shigella* should be considered a single species ^46–48^. Overall, our core genome tree supports previous findings and clearly delineates the genera within *Enterobacteriaceae*.

**Figure 2.**
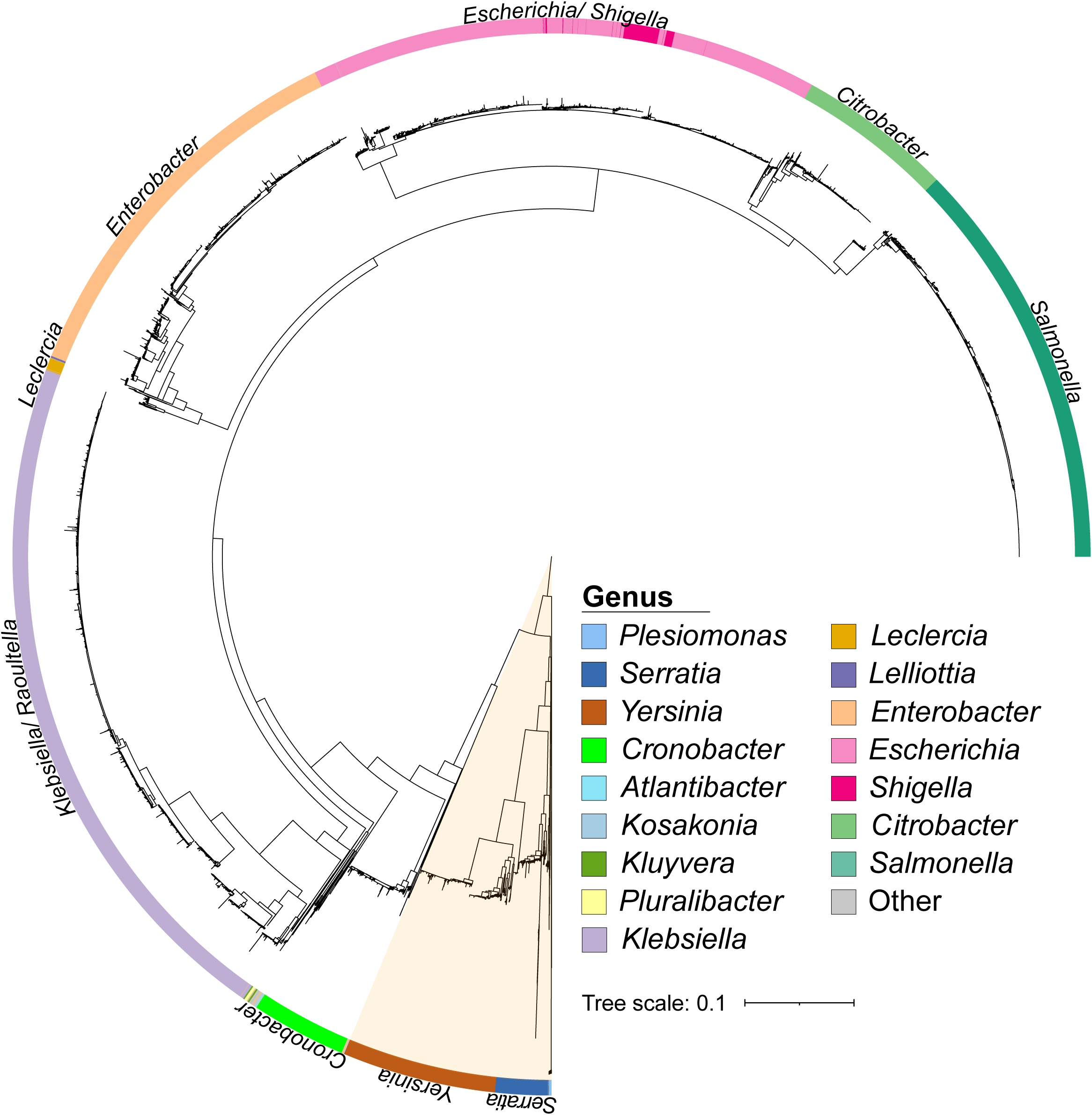
Phylogenetic reconstruction of the *Enterobacteriaceae* family based on 103 concatenated core genes (present in at least 99% of investigated genomes). The tree is midpoint rooted, and the outer ring represented taxonomic classification based on the NCBI taxonomy. Genomes not classified to genus level are labelled as ‘Other’. Peripheral members of *Enterobacteriaceae* (*Yersinia, Serratia and Plesiomonas)* are shaded orange. The concatenated core genes used are listed in **Supplementary Table S2.**

### *Enterobacteriaceae* are adapted to abundant monosaccharides in nature

Our previous phylogenetic analysis validates the broad genome selection for studying the *Enterobacteriaceae* pan-genome. To illustrate the benefit of this approach, we examined the metabolism of the *Enterobacteriaceae* family in depth. Nutrient exploitation is one of the key functions of colonization resistance provided by the microbiome ^17, 49, 50^. Recent publications have highlighted the importance of *Enterobacteriaceae*-*Enterobacteriaceae* competition in effectively limiting or even preventing colonization ^20, 51–56^. This competition is often governed by the differential metabolic utilization of specific carbohydrates. For this reason, we examined the results of our pan-genome analysis to assess energy metabolism, starting with carbohydrate utilization (**Fig. 3**).

**Figure 3.**
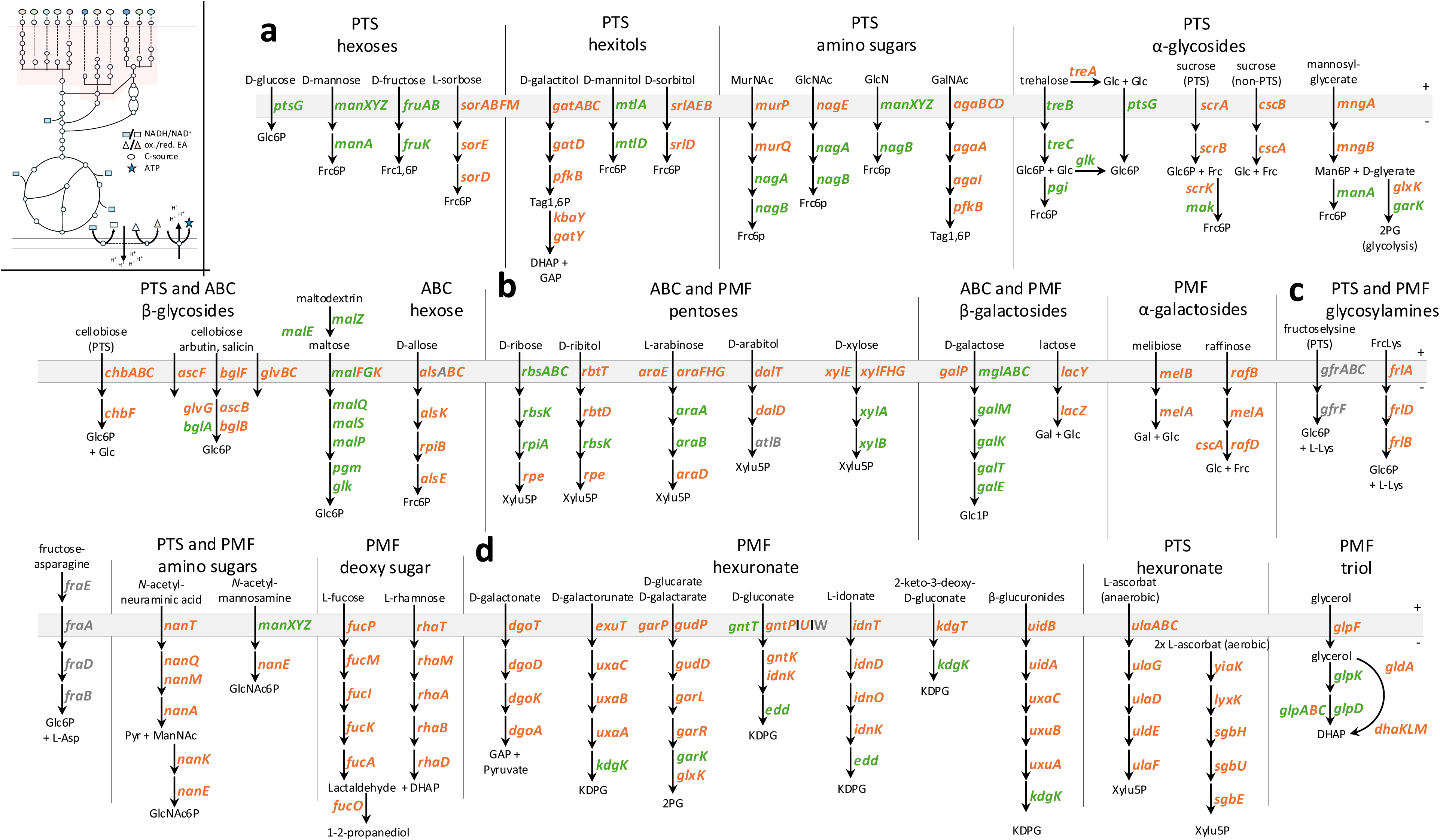
Core and accessory gene distribution of carbohydrate degradation pathways of *Enterobacteriaceae*. A core gene is present in at least 95 % of all analyzed *Enterobacteriaceae* genomes (green). A prevalence of less than 95% is defined as accessory (orange). Genes that were not found in the analysis but have experimental evidence are shown in grey. The top left subpanel indicates energy metabolism, with the respective part of interest marked in red. **a** Depiction of carbohydrates that are almost exclusively transported by the bacterial phosphotransferase system (PTS). Panels (**b, c**) show a mixture of pentoses, hexoses, disaccharides, glycosylamines, and amino sugars that are transported either through primary transport by an ATP-binding cassette (ABC) or via secondary transport through a proton-motive force (PMF)-driven system. **d** Depiction of hexuronate utilization driven by PMF transporters, and the triol glycerol. The different substrate groups are indicated above the metabolic pathways. The abbreviations of carbohydrates are defined in **Supplementary Table S3.**

The main carbohydrate transport systems are the phosphotransferase system (PTS), ATP-binding cassette (ABC), and proton motive force (PMF)-driven transport systems (**Fig. 3**). Carbohydrates transported by the PTS, such as D-glucose, are preferred by *Enterobacteriaceae* ^57^. Genes required for degrading PTS sugars, such as D-glucose, D-mannose, D-fructose, *N*-acetylglucosamine, and D-glucosamine, are found within the core genome (**Fig. 3a, green**). The degradation of hexitols mainly belongs to the accessory genome (**Fig. 3a, orange**), except for D-mannitol (MtlA, MtlD) (**Fig. 3a**). Hexitols and other sugar alcohols present a metabolic challenge under anaerobic conditions of the animal gut, when no external electron acceptor is available, as they generate a reducing equivalent before entering glycolysis. The oxidation of a hydroxyl group (-OH) during sugar cyclization produces an additional reducing equivalent that cannot be disposed of through exclusively mixed acid fermentation. The only disaccharides associated with the core genome are trehalose and maltose, both composed of glucose molecules but differing in their glycosidic linkages: trehalose has a 1,1-linkage, while maltose features a 1,4-linkage (**Fig. 3a**). D-ribose is the only pentose whose transport and degradation belongs to the core genome. Unlike D-ribose, L-arabinose and D-xylose can be transported by two distinct systems: a low-affinity PMF-driven transporter and a high-affinity ABC transporter, both of which are associated with the accessory genome (**Fig. 3b**)^58^. The majority of cytosolic enzymes for degrading L-arabinose and D-xylose are found within the core genome. Thus, L-arabinose and D-xylose utilization is common among *Enterobacteriaceae*, though import occurs via diverse transporters (**Fig. 3b**). Notably, the D-glucose epimer D-galactose is associated with the core genome (**Fig. 3b**). Glycosylamines, which consist of a reducing sugar and an amino moiety, are produced during food processing through the Maillard reaction under heat, forming compounds such as fructose-asparagine and fructose-lysine (**Fig. 3c**). Amino sugars are sugar molecules in which a hydroxyl group is replaced with an amino group, such as D-glucosamine, D-mannosamine, and neuraminic acid. These sugars are also commonly found in their acetylated forms, such as N-acetylglucosamine. Only the pathway for D-glucosamine degradation is fully included in the core genome (**Fig. 3a, c**). Sugar acids, which are hexoses with a carboxyl group (-COOH) at the end of the chain, generally have their degradation genes included in the accessory genome (**Fig. 3d**). Reactive nitrogen species in the gut can oxidize hexoses during post-antibiotic stress, converting D-glucose to D-glucarate, for example ^59^. Additionally, sugars like D-galacturonic acid, D-glucuronic acid, and L-iduronic acid naturally occur in various complex polysaccharides ^60^.

*Enterobacteriaceae* analyzed in this study, share a small number of carbohydrate uptake and degradation systems that are components of the core genome, which are focused on D-glucose, D-glucose epimers, D-glucose-containing disaccharides, and modified D-glucose molecules. This partly mirrors mammalian cells, which have evolved to preferentially utilize glucose ^61^.

### The three glycolytic pathways are associated with the core genome

Most carbohydrate utilization pathways in *Enterobacteriaceae* are linked to the accessory genome (**Fig. 3**). However, the majority of pathways for phosphotransferase system (PTS)-transported carbohydrates - such as D-glucose, D-mannose, D-fructose, D-mannitol, N-acetylglucosamine, and trehalose - that route these carbohydrates into glycolysis are found in the core genome (**Fig. 3a**).

The three glycolytic pathways - namely, glycolysis, the Entner-Doudoroff pathway, and the non-oxidative and oxidative pentose phosphate pathways (PPP)—as well as the tricarboxylic acid (TCA) cycle, are all part of the core genome (**Fig. 4a-e**). Glycolysis, a universal pathway present in both prokaryotes and eukaryotes, metabolizes glucose into two molecules of pyruvate while generating NADH and ATP (**Fig. 4a**)^62^. In contrast, the Entner-Doudoroff pathway in *Enterobacteriaceae* converts hexuronates, while the pentose phosphate pathway processes pentoses into glycolytic intermediates, enabling carbon flux into the TCA cycle (**Fig. 4b, c**). The oxidative pentose phosphate pathway links glycolysis and the Entner-Doudoroff pathway to the non-oxidative pentose phosphate pathway. This pathway provides NADPH for biosynthesis and contributes to the non-oxidative pentose phosphate pathway for biosynthetic precursors (**Fig. 4d**)^63^. The regulatory pathway that governs carbohydrate utilization is the PTS that primarily affects catabolite repression and phosphorylates incoming PTS-transported carbohydrates via a process known as group translocation ^57^ (**Fig. 4f**). In addition, the PTS impacts cyclic AMP-dependent catabolite repression, aligning bacterial proteome allocation with metabolic needs ^64^. The PTS acts as the central regulatory circuit, allowing *Enterobacteriaceae* to optimize growth by adapting to varying carbon sources through a complex hierarchical logic, which has been studied in great detail for *E. coli* K12 ^65^.

**Figure 4.**
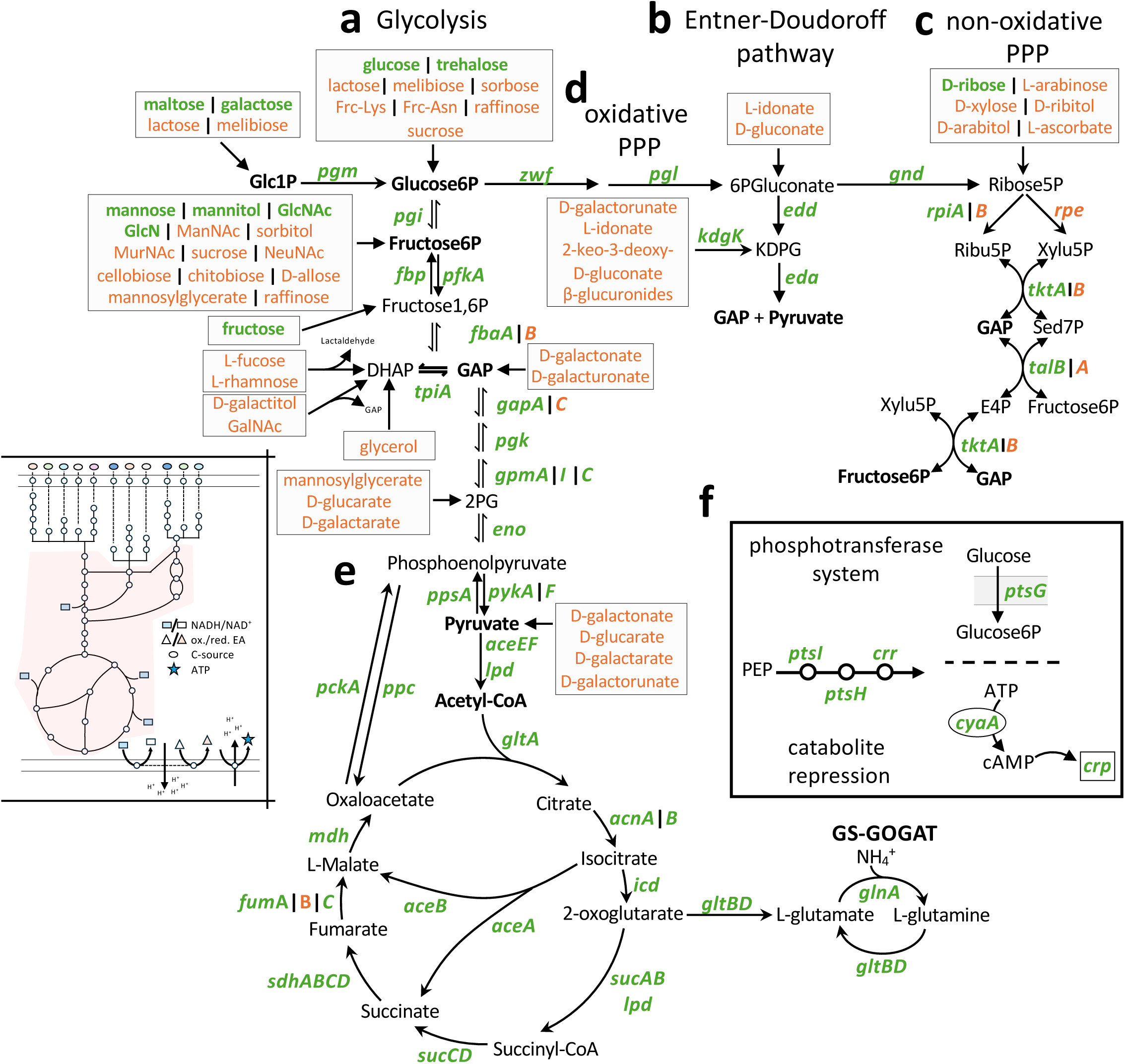
Core and accessory gene distribution of glycolytic pathways, TCA cycle, fermentation, and respiration of *Enterobacteriaceae*. Core genes are defined as present in at least 95 % of all analyzed *Enterobacteriaceae* genomes (green). A prevalence of less than 95% is defined as accessory (orange). Genes that were not found in the analysis but have experimental evidence are shown in black. The top left subpanel indicates energy metabolism, with the respective part of interest marked in red. The three glycolytic pathways— **a** glycolysis, **b** Entner-Doudoroff pathway, and **c** non-oxidative pentose phosphate pathway (PPP)—are shown, including **d** the oxidative PPP, which connects glycolysis to the non-oxidative PPP. The entry points for the various carbohydrate sources from Fig. 3 are indicated by grey boxes. The color-coding shows whether the utilization pathway is part of the core (green) or accessory (orange) genome of *Enterobacteriaceae*. **e** Tricarboxylic acid (TCA) cycle, including the carbon-saving glyoxylate shunt and the glutamine synthetase (GS) – glutamate – 2-oxoglutarate aminotransferase (GOGAT) pathway. **f** Depiction of the phosphotransferase system (PTS), which uses phosphoenolpyruvate (PEP) as a phosphoryl group donor to phosphorylate incoming PTS-transported sugars in a process known as group translocation. Furthermore, the EIIA^Glc^ (*crr*) stimulation of adenylyl cyclase (*cyaA*), which largely controls catabolite repression via cAMP-CRP signaling, is indicated. The abbreviations of carbohydrates are defined in **Supplementary Table S3**.

Interestingly, some isozymes - enzymes that catalyze the same reaction—are linked to the accessory genome, while the primary enzyme is part of the core genome. For example, in *E. coli*, TktA (core) is the main enzyme in the non-oxidative pentose phosphate pathway, while TktB (accessory), an isozyme, has only minor activity ^66^ (**Fig. 4c**). Conversely, *pykF* and *pykA*, encoding pyruvate kinase I and II, respectively, are both associated with the core genome, likely due to their differing physical and chemical properties and kinetic behaviour. For example, PykF is allosterically activated by fructose 1,6-bisphosphate, while PykA is activated by AMP (**Fig. 4a**)^67^. Similarly, fumarases A, B, and C exhibit distinct properties: FumA is a dimeric iron-sulfur cluster-containing hydrolase expressed aerobically and anaerobically, FumB is expressed anaerobically, and FumC is an iron-independent fumarase induced during oxidative stress ^68, 69^. FumA and FumC are part of the core genome, while FumB is not (**Fig. 4e**).

In conclusion, the three glycolytic pathways that provide NADH, NADPH, ATP, and important precursors for biosynthesis are part of the *Enterobacteriaceae* core genome. Isozymes without distinct enzymatic properties are associated to the accessory genome. In contrast to carbohydrate utilization, pathways that route nutrients into the three glycolytic pathways, central carbon metabolism is highly conserved in *Enterobacteriaceae*.

### Respiration of inorganic electron acceptors is associated to the accessory genome

The three glycolytic pathways break down carbohydrates for anaplerosis, supplying important precursors for biosynthetic pathways and generating ATP and NADH for energy metabolism. Energy is conserved in the form of ATP, either by substrate level phosphorylation, or through oxidative phosphorylation. During oxidative phosphorylation, electrons are transferred through a series of redox reactions from a carbon-based electron donor, such as D-glucose, to an organic or inorganic electron acceptor, such as oxygen. Electrons are temporarily stored in the reduced forms of NAD⁺ and the quinones (reduced state: NADH and quinols). Respiratory dehydrogenases, such as *nuo* and *ndh*, oxidize NADH and reduce the quinone pool. In addition to NADH (*nuo* and *ndh*) and succinate (*sdh*) dehydrogenases, which are associated with the core genome, other respiratory dehydrogenases include those for glycerol 3-phosphate (*glpD*), D-lactate (*dld*), L-lactate (*lldD*), D-alanine (*dadA*), pyruvate (*poxB*), and NADPH (*mdaB*) (**Fig. 5a**). Quinols transfer electrons to a terminal reductase that reduces an electron acceptor (**Fig. 5b**). The aerobic (*cyo*) and the microaerobic (*cyd*) oxygen oxidoreductases, as well as the fumarate reductase (*frd*) are the only terminal reductases included in the core genome. The remaining terminal reductases, which utilize inorganic electron acceptors such as nitrate or tetrathionate (S₄O₆²⁻), are part of the accessory genome of *Enterobacteriaceae* (**Fig. 5b**). In brief, this encompasses the electron transport chain, coupled with the translocation of protons to establish a proton motive force, which is utilized, for instance, in ATP production via ATP synthase (oxidative phosphorylation) or in secondary transport ^70^.

**Figure 5.**
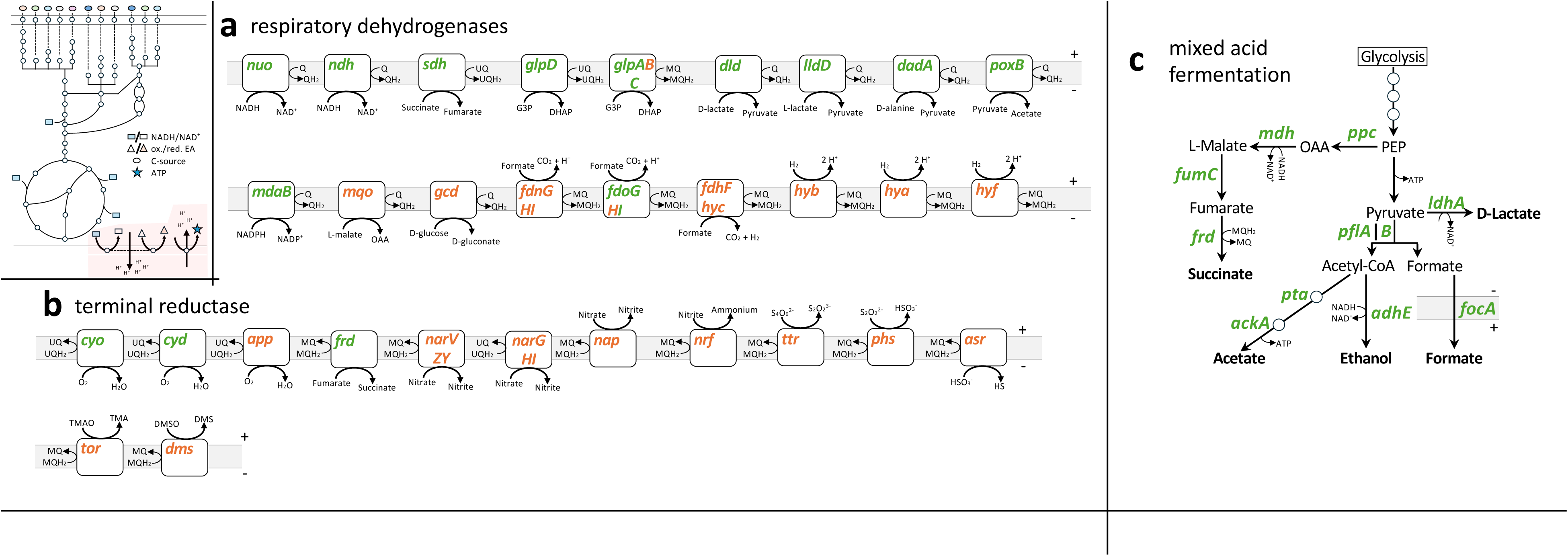
Core and accessory gene distribution of respiratory dehydrogenases, terminal reductases, and mixed acid fermentation *Enterobacteriaceae*. A core gene is present in at least 95 % of all analyzed *Enterobacteriaceae* genomes (green). A prevalence of less than 95% is defined as accessory (orange). The top left subpanel indicates energy metabolism, with the respective part of interest marked in red. **a** Depiction of respiratory dehydrogenases that oxidize an electron donor (e.g., NADH, succinate, etc.) and transfer the electrons into the quinone pool, thereby reducing quinone (UQ and MQ) to quinol (UQH_2_ and MQH_2_). **b** Terminal reductases oxidize quinols and transfer the electrons to an electron acceptor, thereby balancing the redox. The reduced electron acceptor is typically an end product and is secreted. **c** The alternative pathway for redox balancing and ATP formation in the absence of an inorganic electron acceptor is mixed acid fermentation, which exclusively relies on carbon-based electron donors and acceptors for energy and redox balance. The abbreviations of sugars and complete list of individual genes in the respective complexes (e.g., *hyb*) are defined in **Supplementary Table S3 and S4**.

Alternatively, *Enterobacteriaceae* can maintain redox balance and produce ATP during mixed acid fermentation (**Fig. 5c**). This process, a defining feature of facultative anaerobic *Enterobacteriaceae*, produces a range of acidic, reduced end products: succinate, acetate, formate, D-lactate, and the non-acidic ethanol. Acetate fermentation conserves energy in the form of ATP, while succinate, ethanol, and D-lactate fermentation regenerate NAD^+^ and quinones. The acidic and non-acidic end products are exported from the cell, while formate can be used as an electron donor via a respiratory dehydrogenase (*fdn*, *fdo*, and *fdh-hyc*) (**Fig. 5a, c**). The pivotal enzyme in this process is pyruvate formate lyase (PflB), which catalyzes the non-oxidative cleavage of pyruvate into acetyl-CoA and formate. This step is more carbon- and redox-efficient compared to the aerobic counterpart, pyruvate dehydrogenase (*aceEF lpd*), which produces carbon dioxide and NADH. All genes involved in the key steps of mixed acid fermentation belong to the core genome (**Fig. 5c**).

In conclusion, *Enterobacteriaceae*, which are predominantly facultative anaerobic bacteria, possess genes for both aerobic respiration and mixed acid fermentation within their core genome (**Fig. 5**). The distribution of respiratory dehydrogenases and terminal reductases between the core and accessory genomes mirrors the distribution of carbohydrate utilization pathways (**Fig. 3 and 5**). Both types of pathways serve as key interfaces within an ever-changing environment.

### How are the carbohydrate utilization pathways distributed between different strains?

A key component of colonization resistance is nutrient exploitation. This feature is influenced by the complexity of the microbiota and further enhanced by close relatives of the incoming strain, since these will feature a particularly high metabolic resource overlap ^20, 51^. As a proxy for understanding the metabolic resource profiles of *Enterobacteriaceae*, we investigated the presence of different carbohydrate utilization pathways. On average, *Enterobacteriaceae* genomes feature 31 different carbohydrate utilization pathways. The number of detected carbohydrate utilization pathways varied from 3 in a *Serratia symbiotica* strain to 40 in an *Escherichia coli* strain (**Fig. 6a**). We observed a weak correlation between genome size and the size of the metabolic resource profile. For example, symbiotic *Enterobacteriaceae* species exhibit a limited metabolic resource profile, which correlates with their reduced genome size (**Fig. 6b, Supplementary Fig. S1**). To explore the link between microbiota (specifically *Enterobacteriaceae*) complexity and metabolic resource overlap, we assessed the number of strains minimally needed to achieve a high overlap in metabolic resources—defined as 90% of carbohydrate utilization pathways present. When selection was guided by carbohydrate utilization gene analysis (**Fig. 3-5**), a median of just 2 out of 19795 strains sufficed, typically including at least one *Klebsiella* or *Escherichia* strain. In contrast, selecting genomes based on the distribution of *Enterobacteriaceae* in healthy human guts required 7 strains, while selection normalized by the number of genomes per genus required a median of 15 strains (**Fig. 6c**).

**Figure 6.**
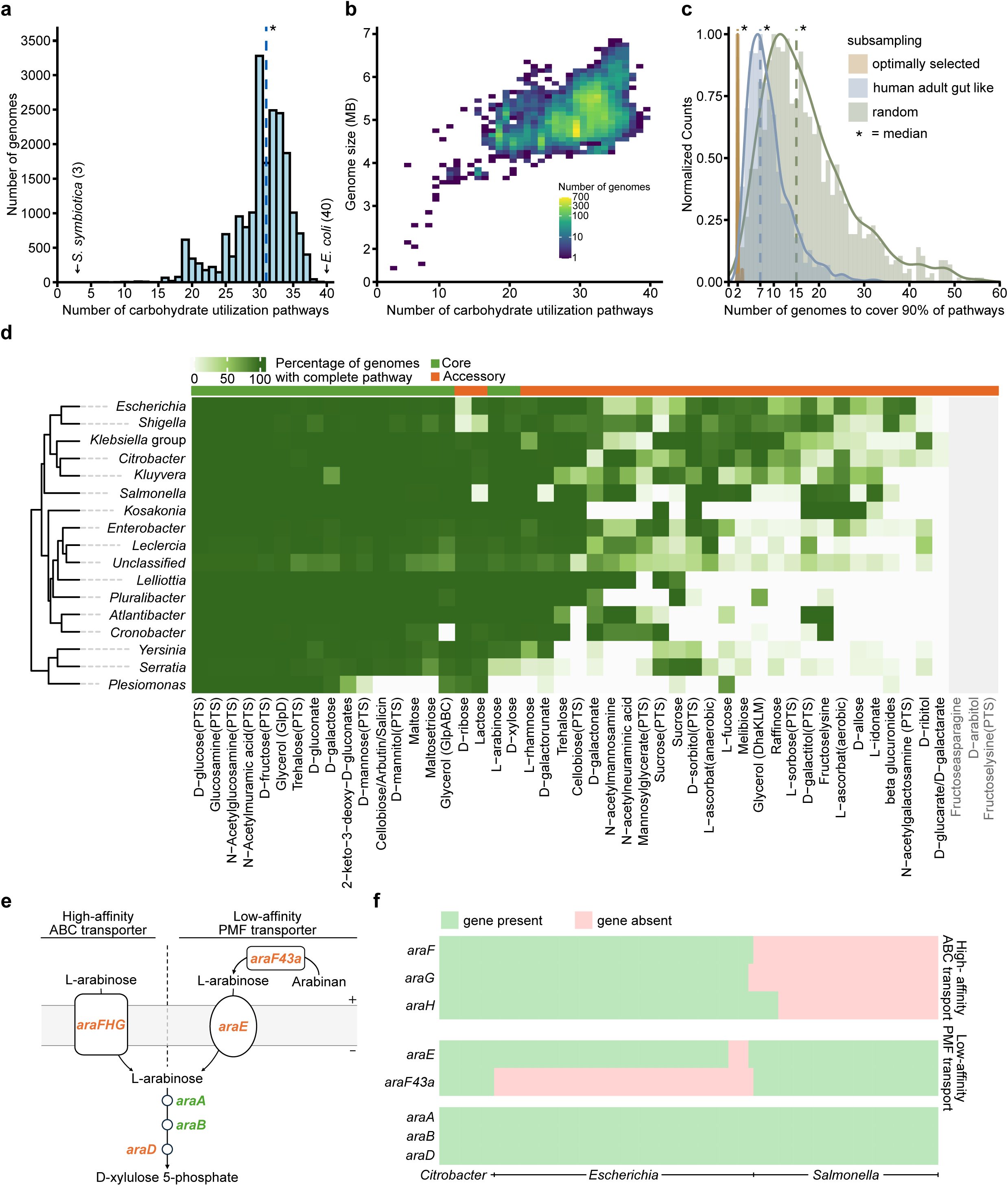
Distribution of carbohydrate utilization pathways in *Enterobacteriaceae*. **a** *Enterobacteriaceae* genomes contain on average 31 carbohydrate utilization pathways (blue dashed line) out of the 49 investigated. **b** The number of carbohydrate utilization pathways is weakly correlated with genome size. Spearman’s correlation: 0.49, P-value < 2.2E-16 **c** Subsampling all investigated genomes while assuming equal probability for each genus, 15 randomly selected *Enterobacteriaceae* strains are on average sufficient to cover 90% of carbohydrate utilization pathways (random, green dashed line). Selecting genomes based on distribution of *Enterobacteriaceae* in a healthy human adult gut, 7 genomes are typically required (human adult gut like, blue dashed line). Constructing a minimal set of genomes to cover 90% of the pathways requires just two genomes on average (optimally selected, orange dashed line). **d** Percentage of genomes containing complete carbohydrate utilization pathways stratified by genera. Rows and columns are hierarchically clustered based on the presented data. Pathways that were not found in the analysis but have experimental evidence are shown in grey. **Supplementary Table S5** provides the full list of genes analyzed for presence in each carbohydrate utilization pathway. Number of genomes per genera are as in Fig. 1b **e** Schematic overview of L-arabinose utilization using low-affinity proton-motive force (PMF)-driven transporter or high-affinity ATP-binding cassette (ABC) transporters. **f** Low-affinity and high-affinity transporters for L-arabinose uptake seem to be mostly mutually exclusive in *Salmonella* and *Escherichia,* whereas both systems are present in *Citrobacter*.

To enhance visualization, we mapped the presence of carbohydrate utilization pathways across all 16 analyzed *Enterobacteriaceae* genera (**Fig. 6d**). **Supplementary Table S5** details the complete list of enzymes included in each carbohydrate degradation pathway assessed. Transporters were generally excluded from this analysis due to their low substrate specificity. Notably, *Klebsiella oxytoca* strains exhibit, on average, the highest number of carbohydrate utilization pathways, which likely contributes to their context-dependent ability to be competitive against *Salmonella* spp. ^20, 53^. Our carbohydrate utilization gene analysis pipeline may help identify the most potent competitors for therapeutic approaches. Additionally, our analysis can shed light on why some *Enterobacteriaceae* encode a low-affinity L-arabinose importer, while others possess a high-affinity version. *Salmonella* strains encode an α-N-arabinofuranosidase, which liberates L-arabinose from arabinan polymers and is important in long-term infection experiments ^71^. L-arabinose can be transported by a low-affinity proton motive force (PMF)-driven transporter and a high-affinity ATP binding cassette (ABC) transporter ^58^ (**Fig. 6e**). Our analysis has revealed that in certain instances, members of the *Enterobacteriaceae* family have evolved to specialize in using one of these transport systems and have consequently lost the other. This observation correlates with the presence of α-N-arabinofuranosidase, indicating that *Enterobacteriaceae* that have acquired this hydrolase are able to locally increase L-arabinose concentrations to an extent that would render a high-affinity transport system dispensable (**Fig. 6f**).

In conclusion, our large-scale pan-genome analysis provides important insights into current questions in the microbiome field. A key goal is the active displacement of unwanted bacteria from the gut through the application of probiotic microbiota consortia ^56^. Understanding metabolic resource overlap and how to rationally design these consortia to maximize this overlap is crucial for displacing problematic strains or using them as prebiotics to enhance colonization resistance against such strains.

## Discussion

Due to the increasing availability of sequencing data, modern bioinformatic analysis has evolved from focusing on single model organisms to encompassing entire bacterial families. This shift allows for a comprehensive exploration of gene distribution, diversity, and genome plasticity. In our approach, we first analyzed a large number of enterobacterial genomes to construct the pan-genome, identifying genes present in over 95% of genomes as core genes, while those found in fewer than 95% are classified as accessory genes ^72, 73^. The *Enterobacteriaceae* family, although not highly abundant in the mammalian gut, plays a disproportionately large role in public health due to some pathogenic members^1^.

Second, this study sheds light on the organization and function of the core and accessory genomes within the *Enterobacteriaceae* family. Notably, we identified a substantial accessory genome comprising non-annotated or hypothetical gene families, even within well-studied organisms like *E. coli*. Remarkably, nearly 50% of *E. coli’s* genome remains either unannotated or only annotated with predicted functions, with around 30% of protein functions still experimentally unstudied ^74, 75^. For less-researched *Enterobacteriaceae* members, this proportion could be even higher. Recent findings in *E. coli* highlight that unknown gene families significantly complicate the pan-genome structure through co-occurrence or avoidance ^76^. We expect similar influences within the *Enterobacteriaceae* pan-genome, where over 70% of gene families are hypothetical. Additionally, the absence of certain pathway components in our dataset may stem from incomplete, inconsistent or missing annotations, as gene annotation continues to present a key challenge in the analysis of prokaryotic genomes and pan-genomes ^31, 77^. Other potential issues in our study may arise from the chosen algorithms for gene family clustering, including the merging of some gene families due to high sequence similarity, unresolved gene fusions, or incomplete fission events. Collectively, these challenges can obscure a comprehensive understanding of pan-genome dynamics.

While our study primarily focuses on the pan-genome of *Enterobacteriaceae* at the family level, the extensive dataset also allows us to explore pan-genome structure at a finer taxonomic resolution. We constructed gene families at various amino acid sequence identity thresholds to accommodate variation in evolutionary rates among gene families ^78^. For more detailed pan-genome analysis within specific *Enterobacteriaceae* species, stricter sequence identity thresholds may be required to account for the narrower taxonomic range. To define species-level pan-genomes fixed thresholds are commonly used in the literature ^27^. Our dataset’s taxonomic classification relies on NCBI annotations, which we validated using 120 marker genes with GTDB-Tk. While genus-level classifications are generally consistent, we observed some discrepancies at the species level, particularly within the genus *Yersinia*, which may affect higher-level analyses. The current pan-genome analysis predominantly focuses on well-studied genera such as *Escherichia*, *Klebsiella/Raoultella,* and *Salmonella* reflecting current sequencing biases. For example over 95% of available *Enterobacteriaceae* genomes in ProGenomes3 originate from these genera ^79^. As metagenomics and the availability of metagenome-assembled genomes (MAGs) continue to advance, we anticipate broadening this focus. This progress will enhance our understanding of gene presences and absences in a population context. Ideally, comparing genomes from various niches will allow us to define enterobacterial ecotypes - strains that use a similar ecological niche -and identify niche-specific genes^80^, providing insights into evolutionary advantages in specific ecological or genetic contexts.

Our study utilized energy metabolism to exemplify how a pan-genome analysis at the family level of *Enterobacteriaceae* enhances our understanding of a key feature of the microbiome: nutrient exploitation. As facultative anaerobes, metabolic genes that enable *Enterobacteriaceae* to thrive in both aerobic and anaerobic environments are part of the core genome. This includes glycolysis, Entner-Doudoroff and the (non)-oxidative pentose phosphate pathway, as well as tricarboxylic acid (TCA) cycle and mixed acid fermentation (**Fig. 4 and 5c**). Noteworthy, most of the pathways required to utilize specific carbohydrates and most terminal reductases are found in the accessory genome, likely reflecting their high context-dependence, i.e. different environmental availability of the respective substrates in different niches. Recent publications have highlighted the availability of carbohydrates in the mammalian intestine, showing that carbohydrates whose degradation belong to the core genome are also abundant in the gut ^81, 82^. This observation largely reflects the sugar monomeric composition of diet, with D-glucose, D-fructose, D-galactose, and D-mannose being the most abundant ^83–85^. Inorganic electron acceptors play a crucial role, particularly during enteric diseases or microbiota disturbances. However, the availability of these molecules is largely context dependent. In the gut, inflammation-induced oxidative stress is necessary to generate nitrate and tetrathionate, which favour the growth of *Enterobacteriaceae* ^86, 87^. At the same time, oxygen becomes available due to the host’s metabolic shift ^88–90^. In contrast, fumarate respiration, a core metabolic pathway, is always used and contributes to *Salmonella* growth in the non-inflamed and inflamed gut because fumarate can be supplied through multiple sources ^91–95^. This is essential because biosynthetic pathways, such as heme *b* and pyrimidine synthesis, rely on fumarate reductase and quinones as electron acceptors under anaerobic conditions. The importance of fumarate respiration is particularly evident in the genera *Escherichia* and *Salmonella* during all stages of infection ^81, 82, 93–96^.

Freter’s nutrient niche theory posits that a bacterial species needs to more efficiently utilize a nutrient niche than the competition ^13, 14^. This theory can translate into differences in bacterial metabolic capacity, such as the presence of genes that allow one species to use a carbohydrate inaccessible to others. Our analysis highlights the high variability in carbohydrate utilization systems among *Enterobacteriaceae* and provides a framework for understanding why different *Enterobacteriaceae* strains can often co-exist in the same intestine ^97^. The sheer abundance of pathways per strain may also explain why *Klebsiella* strains are repeatedly discovered as excellent probiotics for limiting *Salmonella* colonization in certain contexts ^20, 53^. Our pan-genome analysis provides important groundwork for the future design of probiotic consortia aimed at efficient nutrient blocking or exploitation. By maximizing metabolic resource overlap, these consortia could effectively inhibit problematic bacteria that infiltrate a healthy, homeostatic gut microbiome. Interestingly, on average the highest number of carbohydrate utilization pathways were found in a *Escherichia coli* strain, with 40 pathways. It is important to note that defining metabolic capabilities based on bioinformatic analysis has limitations, as the presence of a gene does not necessarily imply the production of a functional protein, as previously demonstrated in *Salmonella enterica* serovars Typhi and Paratyphi A ^98^. No strain possessed all 49 analyzed carbohydrate degradation pathways, raising questions about the cost-benefit balance that governs the number of carbohydrate utilization pathways within each *Enterobacteriaceae* species and why *Klebsiella* harbors so many.

In conclusion, Freter’s nutrient niche theory can be differentiated into shared resources, accessible to various community members, and niche-defining nutrient sources, available to only one particular strain. The former nutrient sources, as evidenced by our pan-genome analysis, will likely include D-glucose, D-glucose epimers, D-glucose-containing disaccharides, and modified D-glucose molecules. In principle, these are nutrient sources that every *Enterobacteriaceae* strain can utilize. Niche-defining nutrient sources are exclusive, accessible by only one bacterial species but not by its competitors; this can provide the decisive advantage for successful colonization. Our carbohydrate-focussed genome analysis strategy may offer a rational approach to select strains for decolonization therapies or mitigating infection risks.

## Acknowledgements

We would like to acknowledge and thank the Hardt and von Mering labs, as well as the NCCR Microbiomes, for their helpful comments and discussions. Many thanks to Yassine Cherrak and Gottfried Unden for providing helpful comments.

## Author contributions

C.v.M, W.-D.H, C.S., L.M. and N.N. conceived and designed the study. N.N. performed the pan-genome analysis. C.S. applied the pan-genome analysis to energy metabolism. N.N. and C.S. wrote the manuscript with contributions from all authors.

## Methods

### Selection of *Enterobacteriaceae* genomes

From the progenomes v3 database ^79^, 525’581 genomes classified as *Enterobacteriaceae* or *Yersinia* according to the NCBI taxonomy were downloaded. First, genomes were screened to select high quality genomes. In a first step, CheckM2 ^99^ assembly statistics, kindly provided by Sebastian Schmidt, were used to remove genomes with completeness below 95% and contamination larger than 10%. Furthermore, highly fragmented genomes (more than 300 contigs) were discarded. Next, genomes with total contig sizes larger or smaller than 1.5 * IQR (inter quantile range) compared to genomes of the same species were removed. Additionally, genomes, with a coding density outside of the 0.5-1.5 range were not considered. Finally, all genomes of species containing less than 10 genomes in the filtered dataset were removed and overrepresented species (mainly *Escherichia coli*, *Salmonella enterica* and *Klebsiella pneumoniae*) were randomly subset to maximally 3200 genomes to generate a *Enterobacteriaceae* genome collection consisting of totally 19’795 genomes. Unless otherwise stated, the NCBI taxonomic classification was used to assign taxonomy to each genome. The identifiers and taxonomic classification of all genomes are provided in **Supplementary Table S1**.

### Pan-genome analysis

All genomes were preprocessed with Prodigal v2.6.3 ^100^ to obtain coding sequences (CDS). We generated a family wide pan-genome by clustering CDS into gene families using PIRATE v1.0.5 using standard parameters and an MCL inflation parameter of 3. Overall, 141’675 gene families were created. Ubiquity was calculated for each gene family and for each genus based on presence and absence. In short, for each genus the number of genomes containing a specific gene family was divided by the total number of genomes for the specific gene family. Taxonomic classification was based on the provided NCBI taxonomy. Gene families were labelled as core if they occurred in at least 95% of all investigated genomes (core genome). All remaining gene families were classified as the accessory genome. Accessory gene families, which were present only in genomes of the same genus were labelled as genus specific, while gene families present in at least 2 genera were labelled as genus unspecific. The core genomes compromised 1’801 gene families (1.3%) while the accessory genome consisted of 139’874 (98.7%) gene families.

We annotated all gene families against the UniProtKB/Swiss-Prot database ^36^ using MMseqs2 v14.7e284 (--start-sens 1 --sens-steps 3 -s 7 –max-accept 10000). Hits with e-value > 1 e-5 were discarded, and the best hit was kept. Annotations were aggregated per gene family and if available, the two most abundant annotations were reported. Sequences which were not annotated were labelled as ‘unknown’. Based on the annotation, gene families were summarized into three annotation status brackets based on the inferred short gene name: Resolved (>= 90% of the sequences contain the same annotation), Ambiguous (< 90% of the sequences contain the same annotation and less than 10% Unknown) and not annotated (>= 10% of the sequences are not annotated). For each gene family a representative sequence was chosen. For annotated gene families, a random sequence from the most common annotation was selected. A random sequence was chosen for non-annotated gene families.

A phylogenetic tree was constructed based on core gene families present in at least 99% of all genomes. Additionally, families with unresolved annotations or families containing non-single copy were removed. The full list is provided in **Supplementary Table S2.** The resulting 103 gene families were subsequently used in the phylogenetic analysis. Briefly, each gene family was individually aligned using muscle v5.1 ^101^ and SNP-sites v2.5.1 ^102^ was used to produce alignments of variant sites. The resulting alignments were concatenated, and a maximum-likelihood tree was constructed using RAxML-NG^103^ with the general time-reversible model GTR+G model with a discrete GAMMA model of rate heterogeneity with 4 categories. ITOL v6.8.1 ^104^ was used to midpoint root the tree and for tree visualizations. Peripheral members of *Enterobacteriaceae* are shaded orange and include *Yersinia*, *Serratia* and *Plesiomonas* ^40^.

### Mapping core and accessory genome on metabolic pathways

The *E. coli* K-12 database EcoCyc.org was utilized as a primary point of reference for metabolic pathways ^105^, and EcoSal Reviews ^70, 106, 107^. The pathways presented in these figures (**Fig. 3-5**) are a curated selection of annotated and referenced research, primarily grounded on *E. coli* and *Salmonella* spp. research. A full list of abbreviations is available in **Supplementary Table S3** Complexes, especially in Fig. 5, are named by their operon rather than individual genes. A complete list of all genes within each complex is provided in **Supplementary Table S4**.

### Investigating the distribution of carbohydrate utilization pathways

We searched the pan-genome for gene families containing annotations presented in **Figure 3**. A complete list can be found in **Supplementary Table S5**. We consider a carbohydrate utilization pathway to be present only if each of its genes can be found within a specific genome. Due to the potential substrate ambiguity of transporters, we only considered intracellular genes, with exception to glucose utilization. Spearman correlation values of genome size against number of carbohydrate utilization pathways present were computed using the cor.test function of stats v4.1.3 in R. The core and accessory status of a carbohydrate utilization pathway is assessed based on the average presence of each gene in all genomes. A pathway is considered ‘core’ if the average is larger or equal to 95%, and considered ‘accessory’ otherwise. Genera and carbohydrate utilization pathways are ordered based on a hierarchical clustering performed using Euclidean distance and complete linkage as implemented in the ComplexHeatmap package in R ^108^.

To calculate the metabolic resource overlap of subsampled genomes, we consider the presence of any alternative pathways for a specific sugar molecule as sufficient to consider this resource as covered, except for L-ascorbat, fructoselysine, glycerol, sucrose and trehalose, resulting in a total of 49 metabolic resources (**Supplementary Table S5**). We distinguish between three different subsampling methods: “optimally selected”, “random” and “human adult gut like”. For the “optimally selected” subsampling, we randomly selected an initial genome as a starting point. Subsequently, the next genomes are chosen to cover most of the missing resources until at least 90% are covered. In random subsampling, we randomly select a genome without replacement at each step until at least 90% of the metabolic resources are covered. To account for uneven distribution of genera in the dataset, each genus was assigned equal selection probability. To subsample according to a human adult gut distribution of *Enterobacteriaceae*, we obtained healthy human gut metagenomic samples from the Unified Human Gastrointestinal Genome (UHGG) v.1.0 catalogue ^109^. To this end, we extracted healthy adult human samples (n = 5128) and read counts of non-*Enterobacteriaceae* species were removed. Additionally, genera not represented in our data set were excluded. Based on this filtered data set, read counts were aggregated by genera and average abundances were calculated by sample. Subsampling was implemented by first selecting a genus based on previously calculated average abundances, followed by randomly selecting a genome from the chosen genus. This process was repeated until at least 90% of the metabolic resources were covered.

To visualize the presence of the high- and low-affinity L-arabinose transport systems and its correlation with the presence of an α-N-arabinofuranosidase, we randomly selected in total 100 *Citrobacter*, *Escherichia* and *Salmonella* genomes. Based on the retrieved annotation, we selected gene families annotated with *araF*, *araG* and *araH* for the high-affinity ABC transport system, *araE* and *araF43*a for the low affinity PMF transporter, and *araA*, *araB* and *araD* for the intracellular pathway. Rare gene families and singletons containing any of the aforementioned annotations are not shown. Genomes and gene families are ordered based on a hierarchical clustering as described previously.

### Data availability

A summary of each gene family created with PIRATE, along with all analyzed *Enterobacteriaceae* genomes and their associated gene family memberships in GFF3 format, are deposited in Zenodo at https://doi.org/10.5281/zenodo.14849847 (ref. ^110^).

## Supplemental Figures

**Figure S1. Symbiotic *Enterobacteriaceae* species exhibit a reduced metabolic resource profile.** Symbiotic *Serratia* species exhibit a reduced metabolic resource profile, measured as number of carbohydrate utilization gene present, as well as a reduced genome size compared to non-symbiotic *Serratia* species.

